# Individual Taste Preferences Predict Cortical Taste Dynamics but Are Modified by Experience

**DOI:** 10.1101/2024.04.02.587752

**Authors:** Kathleen C. Maigler, Jian-You Lin, Ethan Crouse, Bradly T. Stone, Ainsley E. Craddock, Donald B. Katz

**Affiliations:** Neuroscience Program, at Brandeis University, Waltham, MA, USA; Department of Psychology, at Brandeis University, Waltham, MA, USA; Volen National Center for Complex Systems, at Brandeis University, Waltham, MA, USA

## Abstract

As is true in humans, no two rodents prefer precisely the same tastes. Furthermore, no one rodent keeps precisely the same taste preferences forever. Here, we have made use of this between- and within-animal variability to generate and test novel hypotheses about the stability and malleability of neural perceptual coding. We used a brief-access task (BAT) to reveal both individual differences in taste preferences and shifting preferences within individual rats (quantified in terms of lick bout lengths). We moved on to show that these phenomena are not simply random variation: first, by demonstrating that gustatory cortical (GC) taste response dynamics, the late part of which reflect palatability, match that individual rat’s BAT preferences (evaluated almost 2 weeks prior) better than canonical preference patterns; that is, individual differences in preferences reflect differences in neural taste processing. This match, however, links neural taste processing only to the most recent BAT session—GC palatability processing does not reflect performance in earlier BAT sessions. We hypothesized that any tasting experience might impact taste processing (i.e., not just BAT licking), and tested this hypothesis by adding a second session of GC taste-response recordings; palatability-epoch taste responses in this later session no longer matched the most recent pre-recording BAT session, demonstrating that responses following the first electrophysiology session had changed. Together, these data demonstrate that every tasting experience (regardless of the method of taste delivery) changes the rat’s processing of those tastes.

## Introduction

At the most basic level, the task facing an animal’s perceptual systems is to process input such that the animal can generate appropriate behavioral output. This task is of particular importance when said input is gustatory, because taste-guided consumption decisions are made when a potentially dangerous substance is already in the mouth. Palatability (the hedonic value associated with a taste) is the primary variable informing the selection of responses (swallowing or rejection) to such stimuli, and it is therefore unsurprising that a sweet (caloric) taste would be ubiquitously preferred, and a bitter (potentially toxic) taste ubiquitously shunned.

But despite the inescapability of the above logic, individual differences in palatability judgments abound. It is a given, in fact, that different human animals have different specific likes and dislikes when it comes to food; for any particular taste and flavor, there can be found people who love it and people who loathe it. And this is not just true for humans: rat taste preferences vary from strain to strain (Bachmanov et al., 1998; Grill & Bernstein, 1988) and even from individual to individual (Bacharach & Calu, 2019; Inui-Yamamoto et al., 2017; Kotlus & Blizard, 1998; Loney et al., 2012). While part of the explanation for these individual differences is no doubt genetic (Armitage et al., 2025; Bachmanov et al., 2016; Blizard, 1999; Keskitalo et al., 2007), a good deal of evidence supports the hypothesis that eating history controls individual taste preferences. Culture, for instance, drives some of the specifics of food preferences (Jamel, 1996), and simple experience—both social and non-social—is known to impact taste likes and dislikes (Herman & Higgs, 2015; Robinson & Higgs, 2012; Story et al., 2002). Social interactions can change taste palatability for rodents as well (Galef & Wigmore, 1983; Posadas-Andrews & Roper, 1983), and even innocuous experience has an impact on rat taste coding that appears to change preferences (Flores et al., 2016; Flores et al., 2018).

The above work motivates the suggestion that neural taste processing should differ from animal to animal, and even from day to day within single animals, in concert with differences in preference patterns. But there has been very little work done on the relationship between individual differences and neural perceptual responses, and essentially none looking at the stability of this relationship through time. Here we perform this work, repeatedly evaluating rats’ taste preferences using a brief-access task (BAT) that allows rigorous estimates of between-taste preference patterns in a single session, and comparing these behavioral data to assays of the rats’ gustatory cortical (GC) responses to the same tastes—responses that have been reliably and repeatedly shown to reflect palatability in the response epoch spanning the period between ∼0.5s and ∼1.5s following taste delivery (Katz et al., 2001; Mukherjee et al., 2019; Sadacca et al., 2016).

Individual rats’ BAT preferences proved to be a specific match for those rats’ “Late-epoch” GC responses, despite: 1) 2-week “empty stretches” between collection of behavioral and electrophysiological data; 2) a range of sampled taste batteries; and 3) vast differences in taste administration methods used in the two types of sessions (licking in the BAT rig; passive intra-oral cannula delivery in electrophysiology sessions). But this match was delicate, such that even single tasting experiences destroyed it—palatability-related GC responses matched only the most recent BAT preference assessment, not assessments from earlier days, and a second test of the rats’ GC taste responses one day after the first no longer matched the preference ranking from the most recent BAT preference test.

Together, these data demonstrate that: 1) palatability-related spiking in the late “epoch” of a rat’s GC taste responses reflects that individual rat’s taste preferences; 2) these preferences are relatively stable across multiple weeks, provided that the rat does not have intervening taste experience; and 3) the match between a rat’s perceptual and neural preferences vanishes with further taste experience, regardless of whether the experience is self-acquired or experimenter-applied.

## Results

### Experimental Overview

To measure each rat’s taste preferences, we analyzed licking patterns from Brief-Access Task (BAT) data. The BAT provides a rapid and reliable assessment of preferences across an entire taste battery; in particular, analysis of lick microstructure—most notably, calculation of the average size of lick clusters in 10-sec trials—has been shown to be an accurate assessment of the palatability of a proffered taste (Dwyer, 2009; Lin et al., 2012; Lin et al., 2026; Strickland et al., 2018), with smaller average cluster sizes reliably (but see below) equating to lower palatability. Comparison of cluster sizes therefore provided a rigorous, accurate picture of which tastes an animal likes more, and which tastes it likes less (**Figure 1A**). After 3–5 sessions of BAT testing, rats were prepared for subsequent electrophysiology recordings through the implantation of an intra-oral cannula (IOC) and a 32-channel electrode into the primary gustatory cortex (GC; **Figure 1B**). Following recovery, rats received passive taste deliveries via the IOC while GC activity was recorded over two sessions. Following histological examination (**Figure 1C**), only data from rats with accurate electrode placements were included in the final analysis. **Figure 1D** summarizes when each phase of the experiment occurred with a period of approximately 11 days (encompassing surgical recovery and rig habituation) elapsing between the final BAT session and the first electrophysiology recording.

**Figure 1.**
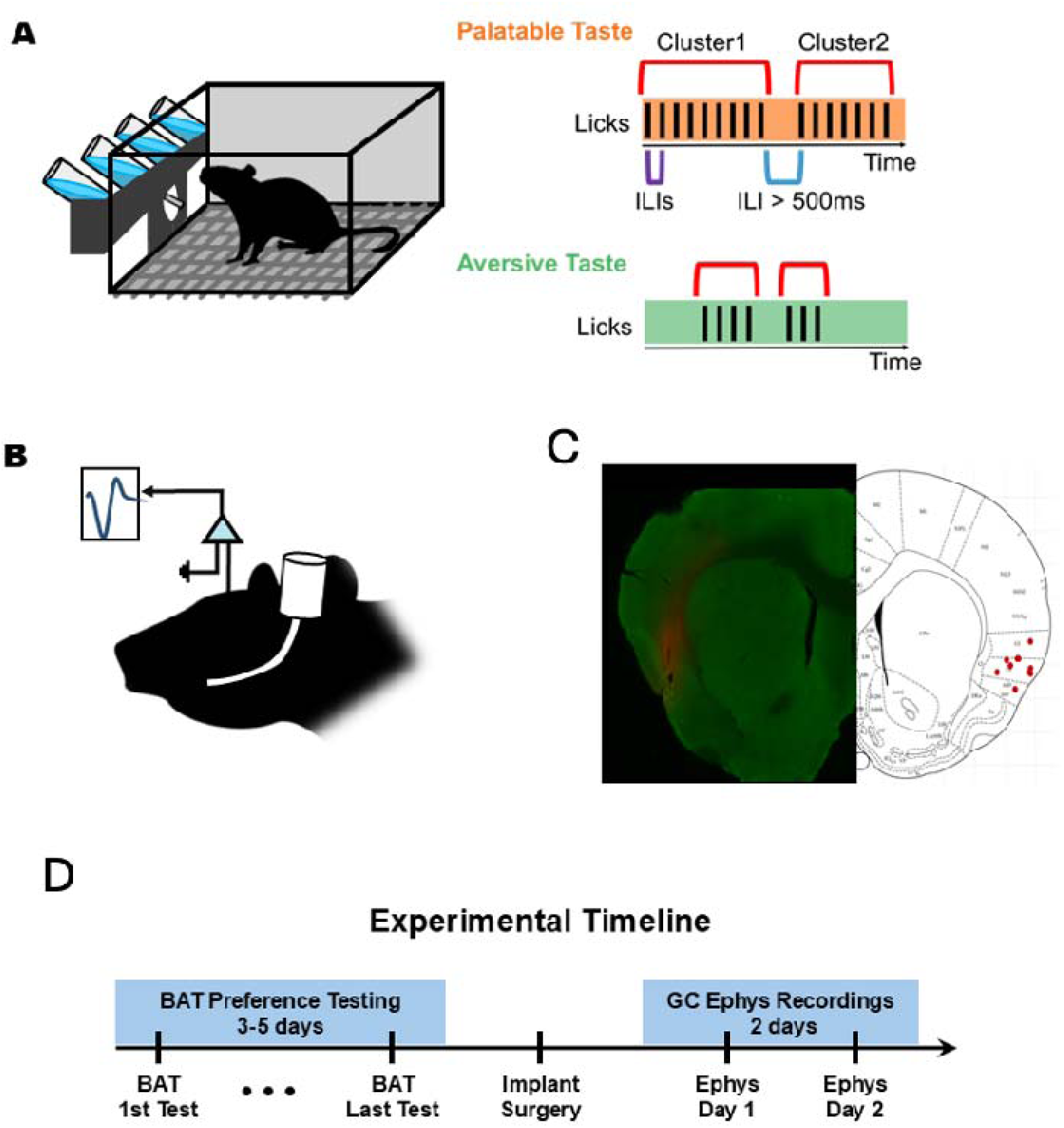
Experimental methods. **(A)** The brief-access rig—multiple bottles, each made available periodically. To the right are example data schematizing the analysis of rhythmic licking to pull out lick numbers and “clusters,” both of which reflect palatability. **(B)** A schematic of in vivo GC recordings with taste delivery via a pre-implanted intra-oral cannula. **(C)** Histological verification of a representative electrode implant site (left) with dye marking the recording site in gustatory cortex. On the right is the schematic of the coronal slice through central GC, with the positions of electrode tips for all animals (N=9) denoted with red circles. **(D)** The experimental timeline, showing preference testing sessions and the delay (for surgery and recovery) between the final BAT day and Electrophysiology recording/passive tastant deliveries.

### Taste preferences vary between animals and across days

**Figure 2A1** shows average sucrose, NaCl, and quinine lick cluster sizes for a pair of representative rats’ initial BAT sessions. While both rats produced very small lick clusters when offered the bitter quinine, and both produced large lick clusters when offered NaCl, they differed notably in their sucrose cluster sizes. A 2-way ANOVA of these data revealed a significant rat x taste interaction (*F*(2,65)=13.44, *p* < 0.001), with one of the two rats preferring sucrose to NaCl (*p* < 0.05) and the other showing little sign even of preference for sucrose over quinine (*p* > 0.05; note that this rat, which was extensively familiarized with the testing context, consumed NaCl and water avidly, suggesting that it simply, if somewhat inexplicably, did not wish to consume the sweet fluid).

**Figure 2.**
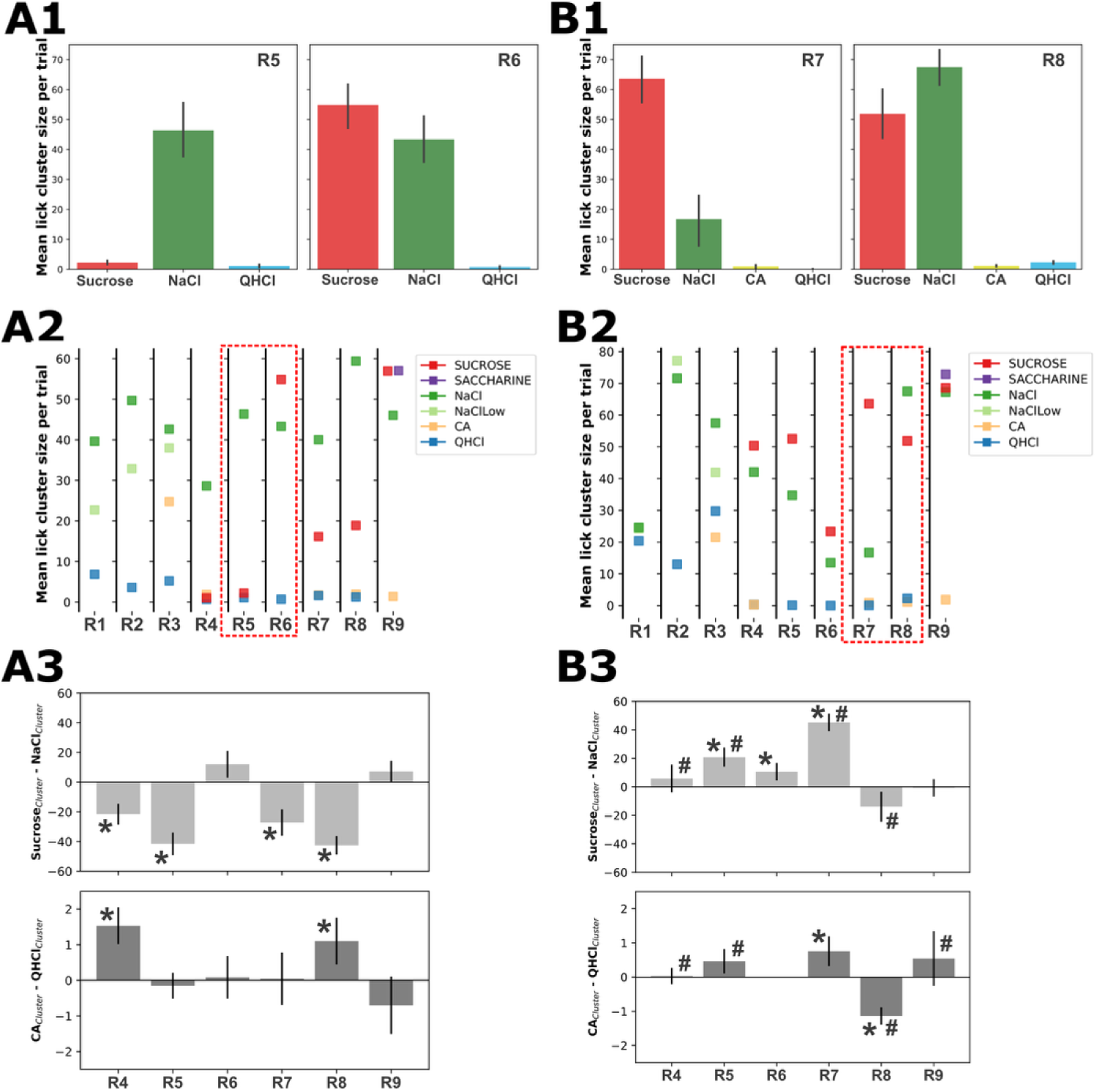
Individual and between-session differences in perceived taste palatability. **(A1-3)** Data from first BAT sessions **(B1-3)** Data from final BAT sessions. **(A1)** Mean lick cluster sizes for two representative rats during the first BAT test illustrate clear individual differences in preference for sucrose, NaCl, and QHCl. A two-way ANOVA revealed a significant Taste × Animal interaction (*F*(2,65) = 13.44, *p* < 0.05). Post hoc Tukey tests further showed that sucrose, while highly preferred by the rat on the left, was perceived similarly to QHCl by the rat on the right (*p* < 0.05). **(A2)** Mean lick cluster sizes for each taste across all animals during the first BAT test (N = 9). The two rats shown in panel **(A1)** are outlined for reference. **(A3)** Differences in lick cluster sizes comparing sucrose and NaCl (top) and comparing citric acid (CA) and quinine (QHCl; bottom) during the first BAT test. One-sample *t*-tests (against a population mean of zero) revealed individual differences in the preferences for the two palatable and the two aversive tastes (*p* < 0.05). **(B1-3)** The same analyses as in A1-3, but for each rat’s final BAT session (depending on the rat, sessions 3 or 5). In **(B3)**, we also show an analysis of whether individual rats changed their relative preferences between BAT sessions—two-way ANOVAs with variables Animal and Session, comparing the data shown in panels **(A3)** and **(B3)**. The majority of rats exhibited significant changes in relative preference for tastes of the same valence across sessions (#, *ps* < 0.05). The labels R1–R9 denote the rats included in the data analysis.

Even casual scrutiny of **Figure 2A2**, which presents lick cluster sizes to every taste across all 9 rats (the red dashed box delineates the rats used for **Figure 2A1**) suggests that while few of the 5 rats that sampled sucrose didn’t like sucrose, individual variability was large enough that individual preference orders often diverged from the canonical palatability ranking (in the case of the battery used with subjects R4, R5, R6, R7, and R8, this canonical ordering was sucrose > NaCl > citric acid > quinine). While the precise battery offered varied from rat to rat, each offered easy canonical ordering (see Table 1 for canonical rankings of each battery), and in many cases actual individual rats’ preferences differed from this ordering.

**Table 1.** Summary of taste stimuli used in brief-access tests (BATs) and electrophysiological recording sessions. NaCl: sodium chloride; QHCl: quinine hydrochloride. The numbers listed after each taste in the Electrophysiology Sessions column indicate canonical rankings (4–3–2–1, from most palatable to most aversive).

|  | BAT Sessions | Electrophysiology Sessions |
| --- | --- | --- |
| **Set 1** | NaCl (0.1M)<br>NaCl (0.05M)<br>Citric Acid (0.009M)<br>QHCl (0.00078M)<br>Citric Acid (0.00625M)<br>QHCl (0.00156M)<br>Water | NaCl (0.1M) ----- 4<br>NaCl (0.05M) ----- 3<br>Citric Acid (0.009M) ----- 2<br>QHCl (0.00078M) ----- 1 |
| **Set 2** | Sucrose (0.3M)<br>NaCl (0.1M)<br>Citric Acid (0.1M)<br>QHCl (0.001M)<br>Water | Sucrose (0.3M) ----- 4<br>NaCl (0.1M) ----- 3<br>Citric Acid (0.1M) ----- 2<br>QHCl (0.001M) ----- 1 |
| **Set 3** | Sucrose (0.3M)<br>Saccharin (0.005M)<br>NaCl (0.1M)<br>Citric Acid (0.1M)<br>QHCl (0.001M)<br>Water | Sucrose (0.3M) ----- 4<br>Saccharin (0.005M) ----- 3<br>NaCl (0.1M) ----- 2<br>Citric Acid (0.1M) ----- 1 |

These differences were easily observed in a comparison of lick cluster sizes to sucrose and NaCl (typically the most palatable tastes) in rats that sampled both (**Figure 2A3-top**), which makes it clear that some rats prefer (this concentration of) NaCl over (this concentration of) sucrose, while others do not. Similarly, differences between citric acid and quinine (both typically thought of as aversive tastes) lick cluster sizes (**Figure 2A3-bottom**) revealed individual differences in how aversive each was to rats that sampled both: some rats disliked quinine more than citric acid whereas several did not. These impressions were statistically confirmed using one-sample t-tests against a null hypothesis of zero difference, with Bonferroni correction applied (*ps* < 0.008; see figure legends for exact *t*- and *p*-values).

Individual differences in taste preferences were not simply a reflection of taste novelty, in that they could also be observed on the rats’ final day of BAT testing (depending on the rat, this was either the 3rd or 5th session; **Figures 2B1-3**—see figure legends for statistics and *p*-values, and note that one rat reliably shunned both quinine and citric acid after an initial lick bout, such that there was no error bar in the equivalent aversiveness of both tastes). These results support the suggestion that perceived palatability varies significantly from rat to rat, even after neophobia—a tendency to avoid novel tastes (Barnett, 1958; Domjan, 1977; Lin et al., 2012) that can itself introduce rat-to-rat differences—has long since subsided.

Direct comparison of **Figures 2A3** and **2B3**, which presents first and final BAT session data from the same 6 rats (out of the total 9 subjects) that sampled both sucrose and NaCl (top panels) as well as quinine and citric acid (bottom panels), makes it clear that rats’ preferences changed with testing: 2-way ANOVAs revealed that the relative individual preference for either sucrose or NaCl was frequently different in the first and final tests (*F*(5,116)=40.28, *p* < 0.001), as was the relative aversiveness of quinine and citric acid (*F*(5,116)=18.34, *p* < 0.001; in both panels, “#” denotes significantly differences between the first and final sessions for that particular rat). That is, rats’ taste preferences changed between testing sessions. Similar changes between first and final sessions were observed in the other 3 rats.

Of course, it is known that experience can specifically increase an animal’s preferences for tastes across time, a phenomenon known as attenuation of neophobia (Lin et al., 2012; Menchén-Márquez et al., 2026; Monk et al., 2014). To determine whether this phenomenon, or any other systematic (i.e., similar across rat) change in taste preference, can explain the changes observed across BAT sessions, we examined the change in lick cluster sizes between the first and final BAT sessions for each taste grouped by canonical “palatable” or “aversive” as above (see Methods). As shown in **Figure 3**, there is no consistent direction in which taste preferences changed for either palatable or unpalatable tastes: some rats’ preferences increased with experience with sampling, others’ decreased; the only reliable effect was change itself. One-sample tests performed on these data revealed that normalized scores—calculated relative to the first BAT session using the formula ∼ Final BAT / (First BAT + Final BAT)—did not differ significantly from 0.5 (i.e., no reliable direction of cross-session change) for either palatable (*t* = 1.91, *p* > 0.05) or aversive tastes (*t* = −0.23, *p* > 0.05); the same results were obtained when analysis was restricted to sweet and/or bitter tastes (sucrose data showed only a nonsignificant trend). In summary, the changes in preference across session were as idiosyncratic as the preferences themselves, which means that attenuation of neophobia cannot account for them (see Discussion).

**Figure 3.**
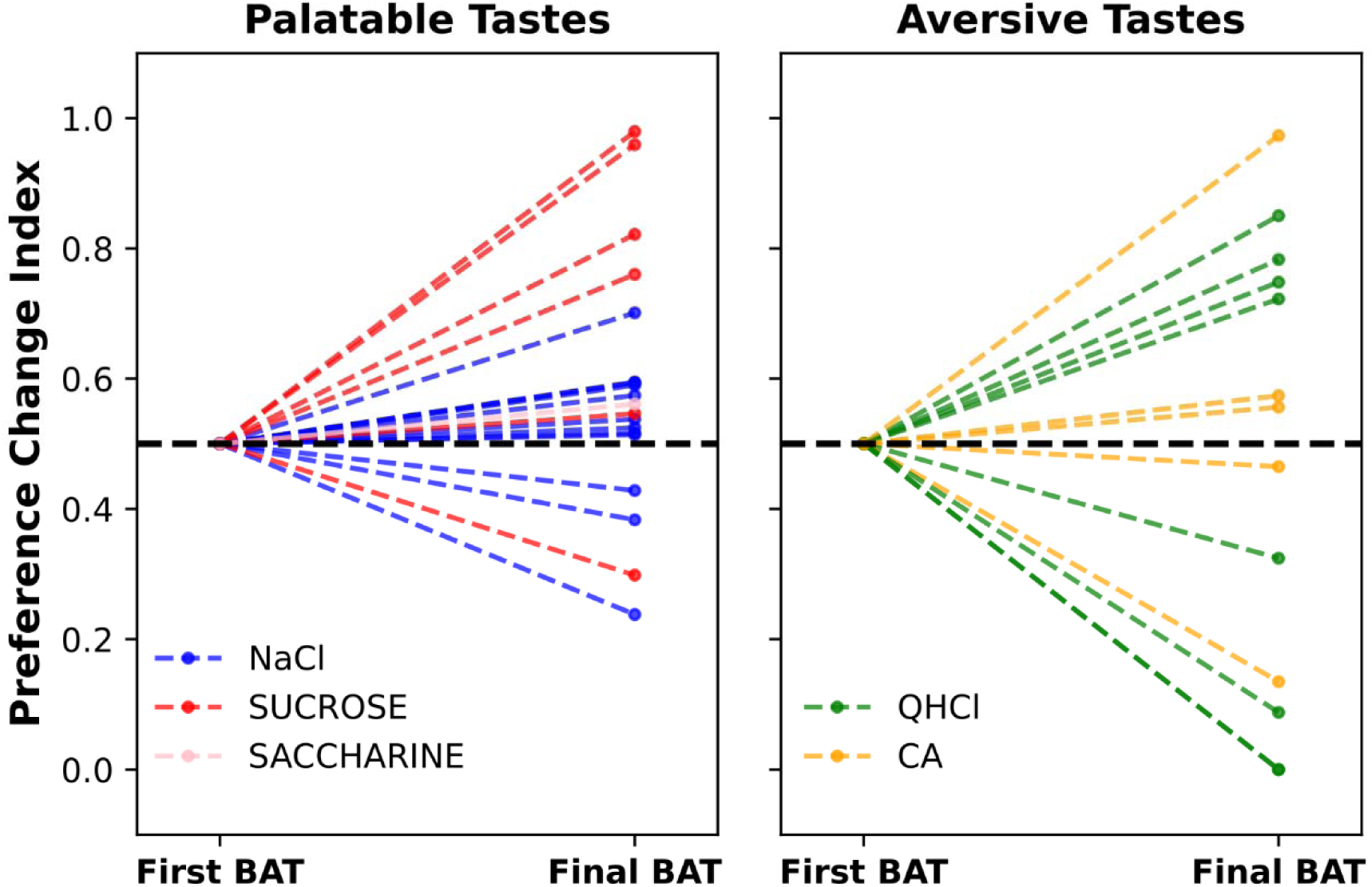
Experience-related changes in taste preferences are not attenuation of neophobia. Each panel shows the change in lick cluster size between the first to final BAT session for either palatable (left) or aversive (right) tastes. Individual tastes are indicated by different colors. In both cases, experience clearly changes behavior, but the changes aren’t in a consistent direction. One-sample t-tests of the preference-change index revealed no significant overall change for either palatable tastes (*t* = 1.91, *p* > .05) or aversive tastes (*t* < 1). The same pattern of results was obtained when the analysis was restricted to sucrose and QHCl. See Methods for details.

### Individual rats’ most recent taste preferences are uniquely reflected in late-epoch GC taste responses

If the between-rat differences measured in the BAT test veridically reflect genuine differences in perceptual preferences (as opposed to representing random variability in behavioral expression of population-general underlying preferences), then they should be reflected in that rat’s neural taste responses. More precisely, the fact that gustatory cortex (GC) activity encodes taste palatability in the late taste-response epoch (Katz et al., 2001; Mukherjee et al., 2019; Sadacca et al., 2016) motivates the specific prediction that responses in this epoch should uniquely correlate with palatability rankings derived from each rat’s own licking behavior (i.e., lick cluster size) during BAT testing.

To test this hypothesis, we recorded single-neuron ensemble spiking activity from GC in the same rats from which we collected preference data, while delivering tastants (the same tastants delivered in that rat’s BAT sessions) through pre-implanted intraoral cannulae (IOCs). A total of 230 GC neurons were recorded (25.5 ±12.1/rat). The bar graphs in **Figure 4A** show the average lick cluster sizes for each taste during first and final BAT sessions from a representative rat: this rat (R3) was exposed to a taste battery including (in order of canonical ranking from most to least palatable) 0.1M NaCl, low-concentration NaCl, citric acid, and quinine; note that the rat’s preferences in the final BAT session differed from this canonical ranking, in that citric acid was deemed least palatable.

**Figure 4.**
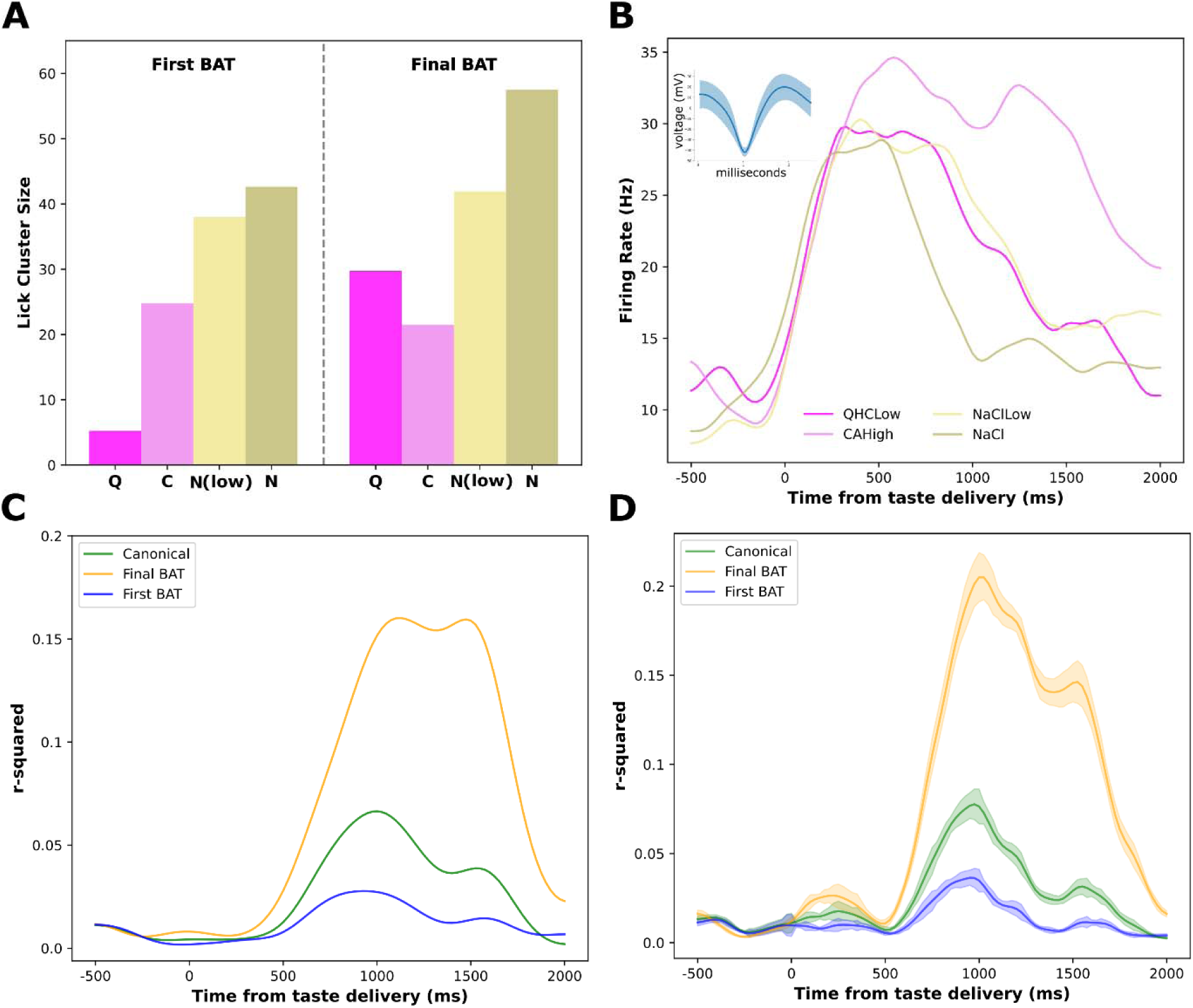
Palatability-relatedness of Late-epoch taste responses match the individuals’ preferences. **(A)** first- and final-session preferences (lick cluster size) for the battery of tastes sampled by one representative rat. The preference pattern changes with experience, and is non-canonical in the final session. **(B)** Peristimulus time histograms for the same tastes, passively delivered through an intra-oral cannula during an electrophysiology session, for a representative cortical neuron in the same rat, showing typical response dynamics including correlation (r-squared, dotted line and right y-axis) to palatability peaking around 1 second from taste delivery. The inset shows example waveforms. **(C)** Palatability correlations from the GC unit in **(B)** calculated based on the canonical ranking or the ranks estimated from the final BAT test. **(D)** The same analysis performed for the entire ensemble of neurons in this example rat.

**Figure 4B** shows overlain peri-stimulus time histograms (PSTHs) for a single GC neuron’s responses from this same rat to the same tastes. The dynamics of these responses replicate those observed in many previous investigations (Katz et al., 2001; Mukherjee et al., 2019; Sadacca et al., 2016): the firing caused by taste administration had little taste-specificity initially, becoming taste-specific only ∼250 ms after delivery; the responses only later came to reflect palatability, as indicated by the appearance of significant correlations between firing rates and palatability after 500 ms of post-delivery time had elapsed (**Figure 4C**). Palatability-relatedness peaked (again consistent with previous reports) at approximately 1 second after taste administration.

Note, however, that the correlation calculated from comparison of neuron firing to the “canonical” palatability rankings (green line) was, while significant, dwarfed by the much stronger correlations between GC late-epoch firing and palatability rankings estimated from the animal’s own lick cluster sizes (NaCl > low NaCl > quinine > citric acid) measured in the most recent BAT (orange line), which reached a peak correlation of ∼0.4 (i.e., an *r^2^* of 0.16). In contrast, when palatability was ranked based on the first BAT session, the resulting correlations were weaker than those based on canonical rankings (blue line). This same pattern observed in this representative neuron was also evident across this one example rat’s full neural ensemble (**Figure 4D**), in which the peak ensemble correlation was approximately 0.45. These results demonstrate that this rat’s GC taste responses were specifically aligned with its own recent preferences.

This match between GC firing and the individual rat’s most recent preference evaluation is particularly striking given the fact that: 1) BAT preferences were obtained almost 2 weeks prior to the electrophysiology session; and 2) the method of taste administration was starkly different in the two sessions—rats obtained their own taste samples through licking in the BAT, and passively accepted IOC deliveries of tastes in electrophysiology sessions. We can therefore conclude that individual differences in taste preferences as measured in the BAT are real, robust to these variables (see Discussion), and stable across weeks of recovery, at least when those weeks contained no additional experiences with strong basic tastes.

But this last point is key, in that sessions of taste experience had a significant impact on preferences—i.e., in that only the most recently assayed BAT preferences were reflected in neural activity. **Figure 5A**, which presents the analysis for the entire dataset (N = 9 rats), confirms that the magnitude of the late-response palatability-relatedness of firing evaluated using individual rats’ final BAT data (orange curve) is more than twice as strong than when using canonical palatabilities (green curve; note that the small overall magnitude of r^2^ values has to do with the lack of data selection—the fact that we did not restrict the analysis to those neurons with late-epoch palatability-related responses), but also reveals that this enhanced match to the rats’ own data is nonexistent when the first BAT session is used (blue curve). The robustness of this difference is underscored by the fact that only 1-3 days separated these different BAT sessions—1 to 3 days of tasting was enough to eliminate the effect. In fact, the peak neural correlation with palatability was slightly (but significantly) higher when canonical palatabilities were used than when earlier BAT sessions were used (we have no explanation for this small effect, but see the Discussion for speculation on the matter). Clearly, repeated BAT sessions significantly altered the rats’ preferences, thereby altering the match to later assays of GC taste responses.

**Figure 5.**
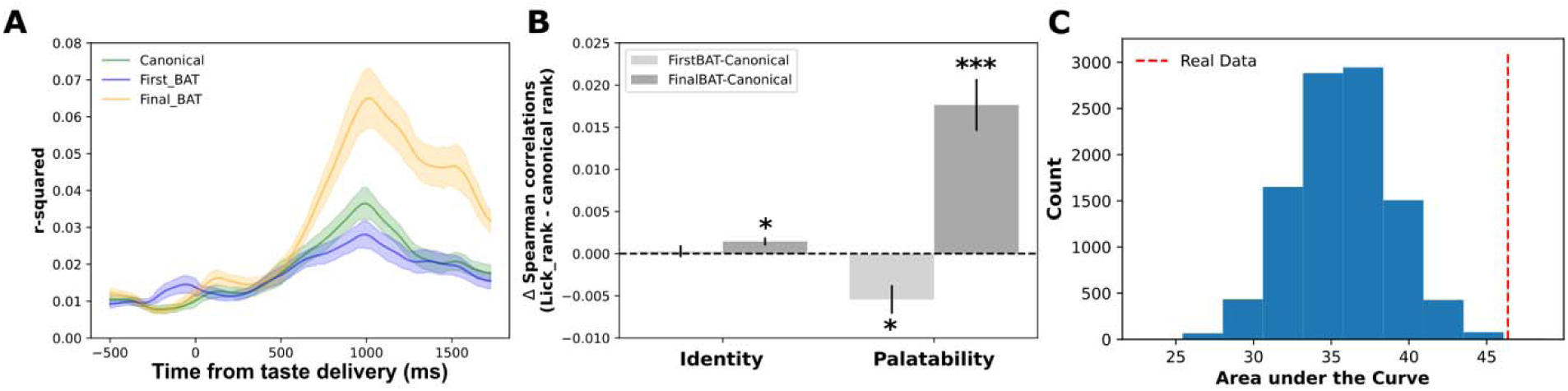
The correlation between Late-Epoch GC firing and an individual rat’s taste preferences is specific to the most recent BAT assay. **(A)**. The palatability correlation over time from all of the (n=129) recorded neurons, using canonical palatability (green), lick cluster data from the first BAT session (blue), and lick cluster data from the final BAT session (orange). **(B)** The difference in correlation calculated from canonical palatability and data from the animal’s individual preferences (First BAT or Final BAT), separated into the Identity epoch (200-700ms) and the Palatability epoch (700-1200ms). * *p* < 0.01, *** < 0.001. **(C)** Magnitudes of the Late-Epoch palatability response (quantified in terms of the area under the curve in the correlation function shown in **5A**) for 10,000 mild shuffles (reversing the order of nearest neighbors—see text for details) of rats’ preference orders. The vertical red dashed line shows the average magnitude for the real data.

**Figure 5B** summarizes these results in a comparison of GC firing and preference patterns, calculated separately for “early” (200–700 ms, a period during which GC responses largely reflect taste identity) and “late” (700–1200 ms, the time during which GC responses are coming to reflect palatability) epoch responses. As expected, effects were minimal during the early epoch, because responses at that time point were not palatability-related; the slight but significant difference between individual preferences measured in the most recent BAT and canonical preferences almost certainly reflects the onset of the Late epoch, a conclusion supported by the lack of significance in a comparison of the effect using first and final BAT rankings (*F*(1,128) = 2.21, *p* > 0.05).

Between-condition differences were pronounced, however, during the late epoch, when GC activity is known to reflect palatability. The brain-behavior correlation was significantly enhanced when using final BAT rankings (compared to when using canonical rankings; *t*(128) = 5.71, *p* < 0.001), and significantly poorer when using first BAT rankings (*t*(128) = −3.20, *p* < 0.01). The difference between these two effects was significant (*F*(1,128) = 30.31, *p* < 0.001). Palatability-related firing, which is restricted to the later aspects of GC taste responses, uniquely reflects the most recent assay of a rat’s individual taste preferences.

As a further test of these results, we performed a control analysis wherein we randomly swap the ranks of “neighboring tastes” of each rat’s final BAT preferences and then generated the plots shown in **Figure 5A** using a random set of that rats’ GC neurons and the sampled preference data. This process was repeated 10,000 times, after which we calculated the area under the curve in the palatability epoch (i.e., the degree to which the correlation rose above baseline across the epoch) to reveal that randomly shifting preferences from the final BAT test made the correlation with GC late-epoch responses reliably worse (*p* < 0.001; **Figure 5C**) These results demonstrate that the match between GC activity and recent preference measurements did not arise randomly, and was specific to the rats’ preferences in that final BAT session.

### Every tasting experience changes GC taste processing

The above results demonstrate that individual differences in taste preferences reflect between-rat variability in neural taste processing, but also demonstrate that tasting experience in the BAT changes those preferences such that there is no match between an individual rat’s palatability-related neural responses and earlier preference tests. An obvious question regarding this result has to do with the specificity of the effect—whether only BAT-type taste experiences (i.e., experiences in which taste samples are acquired *via* licking, under the rat’s volition) alter taste system function in such a way that preferences change. Alternatively, it is possible that any experience with strong basic tastes will alter taste processing.

This speculation engenders a specific novel hypothesis: if only experience with tastes actively licked from a source is sufficient to alter overall preference patterns, then electrophysiology sessions (which involve repeated administrations of each taste to passive rats *via* IOC) should have no such impact; if so, the palatability-related firing observed in a second electrophysiology session should continue to match the most recent prior BAT-based preference evaluation. We tested this novel hypothesis by conducting a second recording session 24 hours after the first to assess whether GC taste responses continued to correlate with licking-based individual preferences (note that between sessions, we advanced the electrode tips by ∼250 µm to ensure that the test was performed on a new set of neurons).

**Figure 6** shows the result of this test. **Figure 6A**, which presents the same moving-window assessment of correlations between neural firing and palatability used in **Figures 4** and **5** (and in many previous publications), once again replicated the late-epoch rise in palatability-relatedness of GC activity, which is robust across different specific measures of taste preference. The neural responses in this second electrophysiology test no longer reflected the rats’ recently-assayed preferences, however, as evidenced by the fact that the correlations with canonical taste preferences, first BAT taste preferences, and final BAT taste preferences were essentially identical—there was no evidence of an enhancement of the GC response correlation with the most recent BAT data when using neurons recorded in the second electrophysiology session. This observation was confirmed by analyses showing no significant difference between correlations with canonical palatability ranks and correlations with ranks derived from either the first or final BAT sessions (*ps* > 0.05; **Figure 6B**).

**Figure 6.**
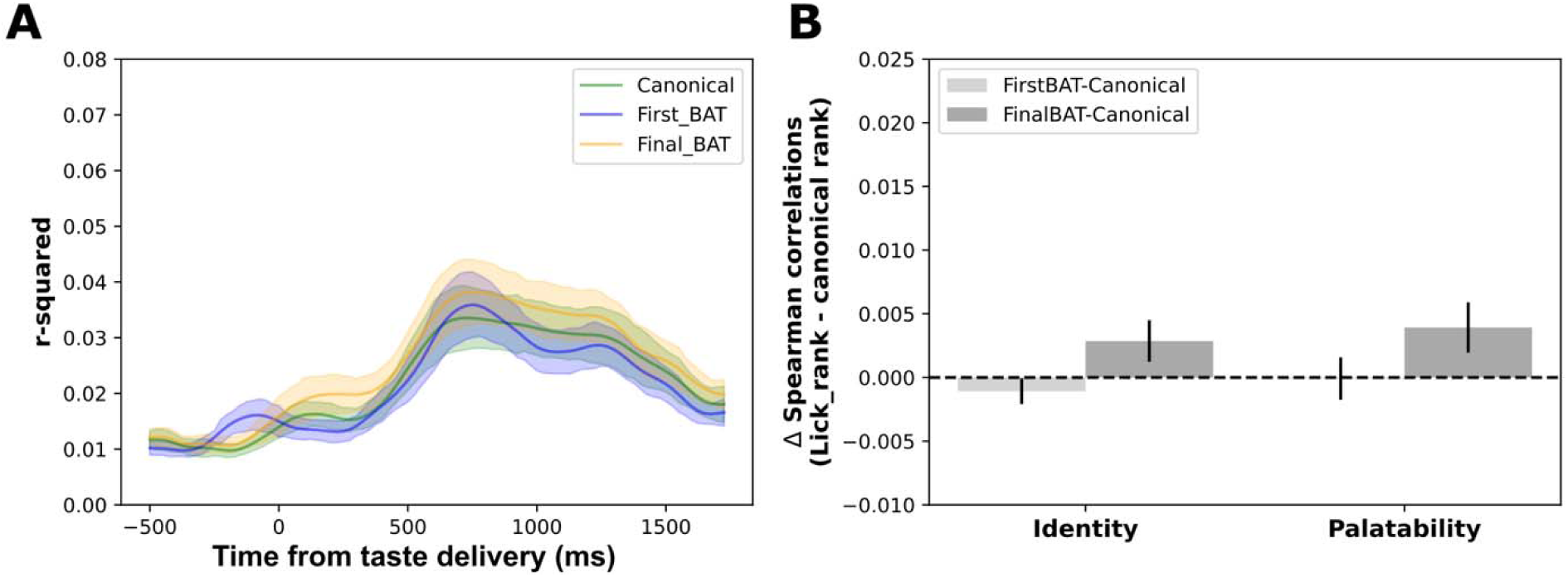
A single session of IOC taste delivery to passive rats nullifies the correlation with palatability calculated from the most recent BAT assay. **(A)** Cortical coding no longer matches rats’ individual preferences when using neurons (n=101) from electrophysiology session 2 (same conventions used in **Figure 5A**. **(B)** A summary of the data in **(A)**, showing there is no longer a significant difference using preference rankings estimated from BAT performance. The Identity epoch and the Palatability epoch activity are taken from 200-700ms and 700-1200ms post-taste delivery, respectively.

While these data do not directly prove that the IOC tasting session (i.e., the electrophysiology session) changed preferences, it demonstrates conclusively that this tasting session changed neural responses so that these responses no longer match the most recently-acquired preference measurements. The most parsimonious conclusion, therefore, is that both lick-acquired and IOC-delivered taste experience quickly changes taste preferences (see Discussion).

We performed further control analyses to test whether the electrophysiology sessions might change some general property of GC responsiveness that could provide an alternative (nuisance) explanation for the reduction in the magnitude of the brain-behavior relationship in the 2^nd^ session. We examined both baseline firing and evoked responses in the identity and palatability epochs (**Figure 7A**), comparing firing magnitudes between sessions. A two-way ANOVA of these data revealed, as expected, that firing rates were higher during responses (Epoch: *F* = 8.26, *p* < 0.05), but revealed neither a Session effect (*F* = 2.06) nor a significant Session × Epoch interaction (*F* = 1.68). Similarly, the number of neurons showing reliable taste responses—defined as significant deviations from baseline activity after taste delivery—did not change from session to session (χ² < 1, *p* > 0.05; **Figure 7B**); nor did the number of palatability-tuned neurons (χ² < 1; **Figure 7C**). As a final check, we looked at the absolute 1^st^-vs-2^nd^ session differences in evoked responses to each taste (**Figure 7D**) and found these differences to be exceedingly small and non-taste-specific (*F* < 1).

**Figure 7.**
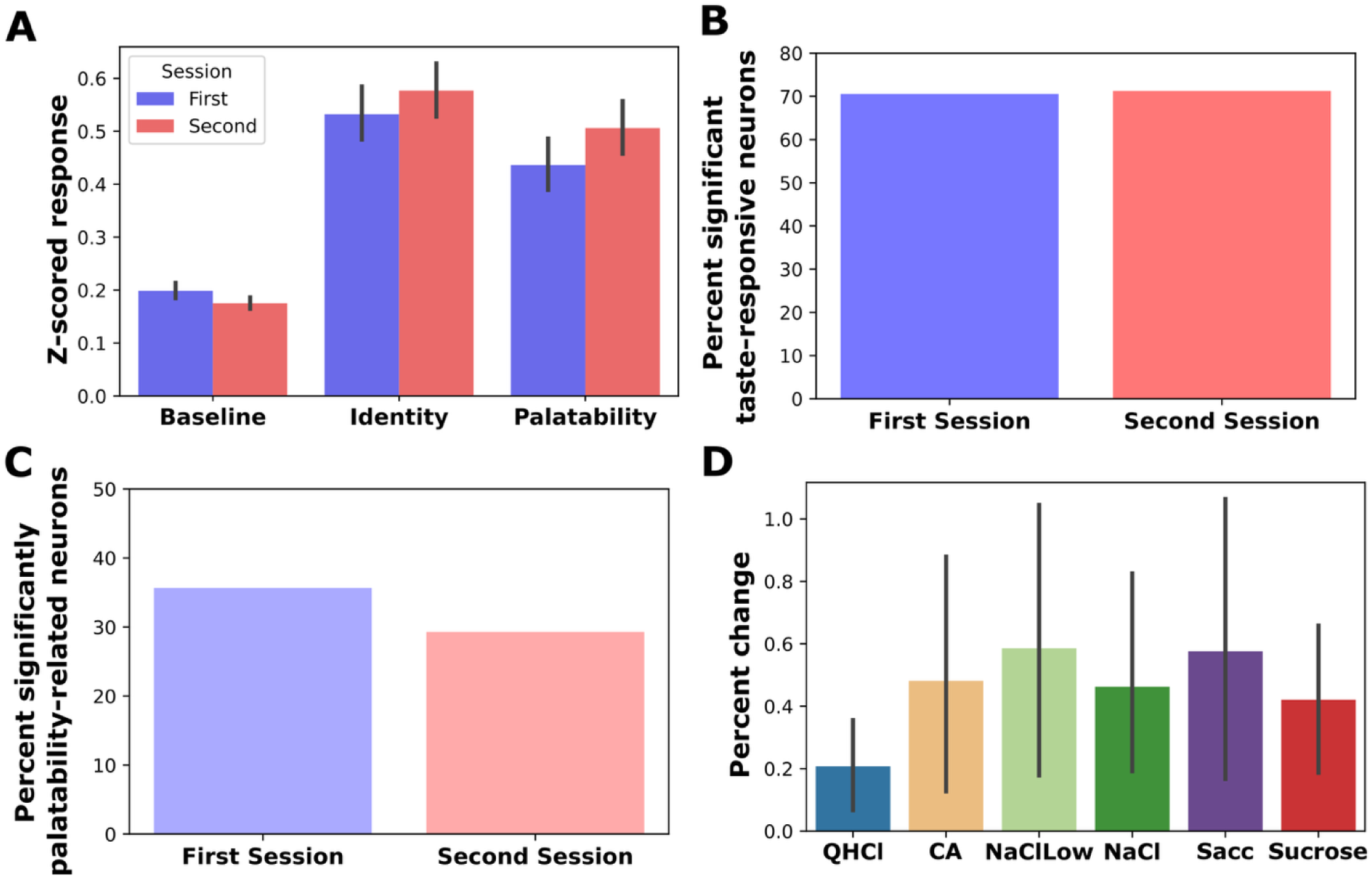
Electrophysiological sessions do not change basic parameters of GC neural activity. **(A)** Normalized activity levels of neurons recorded in the 2 electrophysiology sessions. The first session did not change neuron activity levels. **(B)** The percent of neurons showing significant taste responses is the same for the two sessions (session 1, 70.5% (91 units), session 2, 71.3% (72 units)). **(C)** Similarly, the proportion of neurons producing palatability-related responses in the two sessions did not differ (session 1, 35.7% (46 units), session 2, 29.3% (29 units)). **(D)** The absolute differences in session-1 and session-2 response magnitudes (a number that is necessarily positive) are similarly small—less than 0.6%—for all tastes.

Together, these results indicate that the differential findings between sessions cannot be explained by intrinsic differences in the recorded GC populations, giving us further confidence that what changed with experience was the ordering of the responses in the palatability epoch—that is, the rats’ preferences.

## Discussion

The fact that our BAT data revealed that individual rats have distinct taste preference patterns (i.e., their own ideas of what tastes better and worse) is perhaps not surprising, although the magnitude of those individual differences is almost certainly larger than many would have predicted (particularly given the homogeneity of our rats’ genetics and environment). More surprising is our finding that, even with a surgery, almost 2 weeks of recovery, and a major change in stimulus delivery method separating the BAT and electrophysiological assays, individual behavioral differences are reflected in (“late-epoch”) GC palatability-related spiking. Most surprising, however, is the fact that the stability of perception is upset by taste experience—whether lick-acquired or IOC-delivered, taste delivery had an impact on taste processing. Our results therefore suggest that every new tasting experience changes what a rat thinks of a taste in its mouth; at least as far as gustation is concerned, Heraclitus was correct that “no [animal] ever steps in the same river twice.”

These results validate the use of a brief-access task (BAT), and more specifically lick cluster size analysis, to measure a rodent’s palatability judgments, suggesting that the apparent individual and between-session variability that these tests reveal is *bona fide*. The significantly enhanced match between a specific rat’s measured preferences and their GC late-epoch neural responses (compared to canonical average preference rankings) makes it clear that between-rat variability in BAT performance is not noise but signal—each animal is a genuine individual. An implication of this fact is that any investigation of consumption calculated from BAT data, if rigorously performed, will always have substantial confidence intervals around the group averages. Furthermore, this brain-behavior match provides convergent evidence that the late-epoch GC response, which both predicts and drives the onset of consumption and rejection (Baas-Thomas et al., 2025; Li et al., 2016; Mukherjee et al., 2019; Sadacca et al., 2016), is deeply relevant to behavior; the transition to palatability-related ensemble firing reflects the reaching of a consumption decision, made on the basis of neural processing that is to some degree specific to the animal and session.

This processing turns out to be remarkably stable (note that our point is that the relationships between responses are stable—this does not necessarily mean that the precise responses are themselves unchanging) across days and weeks in which no specific experience with the basic tastes is provided. We performed electrode implantation surgeries after the final BAT preference testing session and gave each rat approximately 11 days to recover prior to the beginning of adaptation to the electrophysiology acquisition setup, such that 2 weeks separated behavioral and neural evaluations of taste palatability. Despite this separation, brain-behavior correlations reflected the individual rat’s BAT performance, a relationship that persisted despite the fact that BAT and electrophysiology sessions differed with regard to: 1) whether the rat was engaged in active sensing (BAT) or passive acceptance (electrophysiology); 2) whether the taste was licked (BAT) or delivered directly into the mouth *via* IOC (electrophysiology); and 3) whether the tastant accumulated in the mouth bit by bit (BAT) or in a single aliquot (electrophysiology). In essence, our experiments constituted a highly conservative test, in that each of the variables discussed above (including the 2-week between-session interval) would be expected to reduce the brain-behavior correlation; the fact that our results were nonetheless highly significant stands as a testament to the robustness of our test and the strength of our conclusions.

Thus, we conclude that “preferences,” as measured by lick bout lengths in the BAT rig, truly reflect the response that the rat would give, were it capable of such linguistic interactions, if asked “what’s the order in which you would rank these tastes in terms of how much you like them?” The BAT data match what’s “in the head”—that is, the content of the specific part of the cortical ensemble taste response that reflects palatability. This interpretation doesn’t invalidate the use of canonical rankings, which clearly provide a first-pass estimate that is significantly correlated, on average, with the rats’ individual preferences. In fact, just as the rat’s idiosyncratic current preferences provide a better match to neural activity than do those canonical rankings, the “incorrect” idiosyncratic BAT data could be expected to match GC responses less well than canonical rankings. We suspect that this is the explanation for the relatively low correlation between GC correlations and non-current preferences shown in **Figure 5**.

Regardless, this last fact also drives home the fact that preferences prove delicate and dependent upon a lack of experience with sapid tastants between the BAT and recording sessions (note that in their home cages, rats may choose to partake only of bland chow and highly familiar biological detritus, and have no access to any strong exemplars of the basic tastes). Rats showed robust changes in their preferences across multiple BAT sessions, and only the preference pattern from the final BAT session matched the palatability-related GC firing recorded during the subsequent IOC session. These results suggest that taste preferences are not fixed properties but are instead updated following each taste experience.

Furthermore, it is clear that even delivering tastes *via* IOC to passively tasting rats has an impact on taste processing, in that the alignment between recorded GC activity and previously estimated preferences is lost in a 2^nd^ passive tasting session. We would argue that the most parsimonious interpretation of these results is that even passive taste exposure via IOC is sufficient to modify a rat’s preference structure, although this was not directly tested here. Fully supporting this conclusion will require a future study in which we incorporate an additional BAT session immediately after the first recording session to test whether this most recent estimate of taste preference best predicts subsequently recorded GC activity. It would also be important to bring the analysis down to a truly rat-by-rat level, which we were unable to do with the current (too-small) dataset, in order to test whether individual animals with non-canonical preferences disproportionately contribute to the observed effects and to determine how individual behavioral preferences relate to neural responses across time.

It is already well known that a first experience with a particular taste can increase the palatability of that taste—a phenomenon known as the attenuation of neophobia (Lin et al., 2012; Menchén-Márquez et al., 2026)—and change the processing of other tastes (Flores et al., 2016, 2022). However, the phenomenon characterized in this current dataset is almost certainly not related to neophobia, in that: 1) the changes observed here occur with each new session, even when the tastes can no longer be thought of as novel; and 2) our analyses demonstrate that these session-to-session changes cannot be described simply as an increase in each taste’s palatability (also, had the palatabilities of all tastes increased together, preference ranks would have remained unchanged). Rather, our data suggest that aspects of taste processing that can otherwise remain stable across at least a 2-week period are altered by each and any taste experience, even after many taste experiences have already taken place.

Clearly, the specifics of taste processing are sensitive to external factors—tastes must be presented to change taste preference structure—but it seems that a range of types of experiences, in which the rat is offered any of a number of different possible taste batteries, are sufficient to evince such changes. It’s worth noting that our rats were housed and tested in a controlled environment (home cages and behavioral rigs); this fact, and the seeming randomness of changes occurring with tasting experience (see **Figure 3D**), makes it reasonable to conclude that the individual differences observed here are likely driven primarily, if not entirely, by idiosyncratic internal factors such as genetic and/or developmental variability, or perhaps lasting impacts of early-life taste experience (see Schiff et al., 2023). Regardless of the source of these individual differences, GC activity reflects palatability judgments of currently experienced tastes, and these judgments vary between rat.

By taking advantage of the naturally occurring individual differences in taste preference, the present findings demonstrate that both the stability and plasticity of these preferences are reflected in the aspect of GC activity that predicts and drives consummatory behavior. In the absence of taste stimulation, taste preferences remain largely stable. In contrast, whenever taste experience occurs, preferences change accordingly. This coordinated change between brain and behavior supports the idea that GC’s role in taste processing extends beyond simple sensory representation: it also contributes to the modulation of consummatory decisions by integrating sensory input with contextual and experiential information. In other words, GC functions not merely as a sensory encoder or decoder, but as an integrative hub that links perception with consumption behavior.

## Materials and Methods

### Subjects

Adult female Long Evans rats (N = 9; 250-375g at time of surgery) were acquired from Charles River Laboratories (Wilmington, MA), singly-housed in independently-ventilated cages on a 12h/12h light/dark schedule, and acclimated to the facility and handling for 1 week before the start of experimental procedures. Unless otherwise specified, animals in home cages had ad libitum access to lab chow and water except where noted. All procedures were conducted in accordance with the guidelines established by the Brandeis University Institutional Animal Care and Use Committee.

### Brief Access Task (BAT) Preference Testing

The BAT rig (Davis MS-160 Lickometer apparatus, Med-Associates Instruments) assessed individual preferences for each taste in the battery. Before preference testing, rats underwent three days of 45-minute habituation to the experimental chamber, after which they were placed on water restriction and given two days of licking training—60 10s-trials in which water was available *via* a metal lick spout attached to glass bottles.

For the preference test assessment, rats were presented with a battery of tastes (in each rat, one of 3 different possible batteries of tastes were used to ensure that results were generalizable—see **Table 1**; as no differences in effects were observed between taste batteries, data was collapsed in our analyses) that spanned a putative range of palatabilities. One taste was offered per trial, in which rats were given 60s maximum to instigate licking, a maximum access time of 10s following the first lick, and a 30s inter-trial interval between trials. Tastes were presented in a pseudo-randomized order (i.e., blocking design), such that a single tastant cannot be offered more than two successive trials (which would adversely impact willingness to participate). Each tastant was presented 8-10 times, depending on the specific taste battery. BAT testing persisted for 3 (taste set 1 & 2) or 5 (set 3) consecutive days.

Tastes, which were dissolved in deionized water (Millipore Milli-Q water purification system), were purchased from ThermoFisher Scientific at ACS Grade (with the exception of saccharin, which was FCC/USP Grade). A fan above the rig served to reduce any potential odor cues that might have accompanied taste presentation. The time of each lick was recorded for off-line preference analysis. All experiments were run between 10:00-16:00.

### Surgical implantation of GC electrode and Intra-Oral Cannula

After the final BAT session, animals were removed from water deprivation and allowed to recover to full, pre-restriction weight for implantation surgery. Rats were anesthetized with an intraperitoneal injection of ketamine (100 mg/kg) and xylazine (5 mg/kg), and craniotomies were made to expose GC. The drivable electrode, consisting of a bundle of 32 nichrome wires (25 μm in diameter), was implanted just above GC (A/P = +1.4, M/L = ±5.0, D/V = −4.4), and secured in place with dental acrylic. Rats also received an intra-oral cannula (IOC)—a polyethylene cannula inserted behind the maxillary molars, through the left masseter muscle and through the opening of the scalp.

Rats recovered for 7 days; during the first day, meloxicam (Alloxate, 5mg/ml, Patterson) (1mg/kg) was injected for pain management.

### Electrophysiological recording with passive tastant delivery

After recovery from surgery, rats habituated for 2 days (30-minute sessions) to the electrophysiological recording rig (plugged into their electrode harness in a plexiglass experimental chamber measuring 8.5 x 9.5 x 11.5 in), after which they were placed on water restriction in preparation for recording experiments. Experiments began with two days of habituation to liquids delivered through the IOC, during which 60 (day 1) and 120 (day 2) trials of water were delivered with inter-trial intervals of 22s. Twenty-four hours after, each rat received two consecutive recording days where 30 randomized presentations of each tastant (a total of four tastes selected from the same taste set received during the BAT preference testing; **Table 1**) were delivered.

Delivery of each taste was individually controlled by solenoid valves, pressurized by inert nitrogen. To ensure our ability to deliver precise amounts of fluid onto the tongue of each IOC delivery, we calibrated the rig to deliver 30 microliters in each aliquot at the beginning of each recording session, controlling the open time of valves *via* a Raspberry-Pi microcomputer.

Spiking data were collected *via* Intan RHD2132 analog-to-digital chip amplifier. Recordings were taken at a sampling rate of 30 kHz. Electrodes were driven down (∼0.25mm) 24hr prior to each recording session to capture different populations of GC neurons for each session. The entire recording rig was enclosed in a faraday cage to reduce latent electromagnetic interference during recordings.

### Electrophysiology data processing

Bandpass filtering and common average referencing were used to clean data and increase signal quality. Discriminable action potentials of no less than 3:1 signal-to-noise ratio were isolated online from each signal and saved digitally. Prior to analysis of taste-related activity, cells were clustered using 3-D cutting techniques alongside supervised verification of inter-spike interval plots. All spikes included in analysis had at least 2000 waveforms and met our criteria of < 0.5% 1ms violations and <2% 2ms inter-spike interval violations. For a comprehensive explanation of the protocol used for spike sorting, refer to Mukherjee et al. (2017). By driving down the mini-microdrive between sessions, new cells were captured each day, which was verified by comparing waveform shapes.

Semi-processed spiking arrays were then exported to an analysis framework built in python. After processing, each 7000ms (2s pre-stimulus, 5s post-stimulus) recording window was binned using a sliding window average with a window size of 250ms and a step size of 25ms.

### Data analysis

#### Computational analysis of BAT data and preference ranks

Lick data were processed and analyzed in custom Python scripts. Libraries integral to the analysis included: pandas, numpy, pingouin, matplotlib, and seaborn. Several lick parameters were measured for each tastant presentation; we used the inter-lick interval (ILI; ∼125msec) to determine lick cluster size—the number of licks produced prior to a 500ms pause (**Figure 1A**). Individual taste preferences were estimated using averaged lick cluster sizes obtained from a BAT session. We used lick cluster size as a proxy for taste preference because this measure has been shown to reliably estimate taste palatability during free licking. Furthermore, we verified that, similar to average licks per trial, lick cluster size derived from brief licking trials (e.g., 10-s BAT trials in the current experiments) yields palatability estimates that are comparable to—and less noisy than—those obtained from the more traditional 15-min ad libitum licking sessions (Lin, et al., 2026).

A similar result was observed when using the first lick cluster, rather than averaging clusters within a trial, to estimate taste palatability. This finding further supports the reliability of the measure (i.e., average lick cluster size), even in some cases when clusters are truncated at the end of a trial.

#### Assessment of preference changes across repeated BAT sessions

Taste preferences are not necessarily constant and can be altered by experience (Lin et al., 2012; Menchén-Márquez et al., 2026; Monk et al., 2014). To determine whether repeated taste consumption across BAT sessions altered taste preference, we compared the average lick cluster size (used as a proxy for taste preference) for each taste between the first and final BAT sessions. To quantify changes across sessions, we treated the lick cluster size from the first BAT session as the baseline and calculated a preference-change index using the following formula:

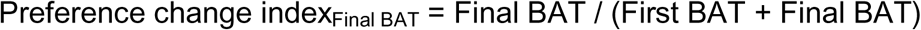

This index captures both the direction and relative magnitude of changes in preference across BAT sessions. A value of 0.5 indicates no change in preference between the first and final sessions. Values >0.5 indicate an increase in preference, whereas values <0.5 indicate a decrease in preference. To determine whether there was a significant systematic change in preference across sessions, we performed one-sample *t*-tests to test whether the mean preference-change index differed from 0.5, the value expected in the absence of a change in preference.

#### Palatability correlation

The method of calculating correlation between GC activity and palatability has been described previously (Katz et al., 2001; Sadacca et al., 2016): a vector of firing rates in each taste response at each 25-ms timepoint in a moving average is Spearman correlated (to control for nonhomogeneity) with the vector of preferences. The canonical rankings for taste batteries are defined and labeled in **Table 1** from most to least palatable based on our BAT experiments and those of others (Sadacca et al., 2012; Spector et al., 1993), which have shown that 0.1 M NaCl is more palatable than 0.05 M NaCl. To incorporate individual preferences here, we also correlated neural activity to animal-matched mean cluster values in the first or most recent BAT session (First BAT, Final BAT).

#### Taste responsiveness and neural excitability

Firing rates were z-scored to the pre-stimulus period by neuron and the mean z-scored firing rate was extracted for each epoch (500ms, baseline: −500:0ms) and compared using a mixed two-way ANOVA (Session × Epoch).

Neurons were categorized as taste responsive if the evoked response (0-2000ms post-stimulus delivery) significantly deviated from pre-stimulus baseline firing (−2000ms-0ms) based on a paired t-test. To qualify as palatability-related, Spearman correlation values to the Final BAT session values had to achieve significance (*p* < 0.01) for ≥3 consecutive bins within the palatability epoch. The proportion of palatability-related neurons per session was compared using χ² contingency tests.

To quantify absolute changes in evoked responses across sessions for each taste, we computed the absolute difference in mean evoked firing (0–2000ms) for each taste between Session 1 and Session 2 and reported as the percent change. These values were compared using a one-way ANOVA across tastes. Significance was set at alpha < 0.5 unless indicated otherwise.

### Histology

Following the experimental sessions, subjects were deeply anesthetized with ketamine/xylazine mix (200, 20 mg/kg respectively, delivered *via* intraperitoneal injection) and perfused *via* intra-ventricular perfusion of saline, followed by 10% formalin. Brains were then extracted for histological verification of electrode placement. Electrode cannulae were painted with a fluorescent cell-labeling dye (Vybrant DiI, invitrogen) upon insertion to verify electrode location and regional markers were used to confirm the cannula track is above and the electrode tip sits in GC as confirmed by another independent experimenter. Following histological examination, one of the 10 experimental rats was removed from the experiment due to electrode misplacement, which left a total of 9 rats in the dataset (**Figure 1**)

## Data availability

The data within this study can be made available from the corresponding author upon reasonable request. Spike sorting code used here as described above is available at https://github.com/narendramukherjee/blech_clust. This study’s design and analyses were not pre-registered.

## Acknowledgements

This work has been supported by National Institute on Deafness and Other Communication Disorders Grants R01-DC006666 & DC007703 to DBK and F31 - DC019863 to KCM.

